# D2 dopamine receptor expression, sensitivity to rewards, and reinforcement learning in a complex value-based decision-making task

**DOI:** 10.1101/2022.02.18.481052

**Authors:** Cristina Banuelos, Kasey Creswell, Catherine Walsh, Stephen B. Manuck, Peter J. Gianaros, Timothy Verstynen

## Abstract

In the basal ganglia, different dopamine subtypes have opposing dynamics at post-synaptic receptors, with the ratio of D1 to D2 receptors determining the relative sensitivity to gains and losses, respectively, during value-based learning. This effective sensitivity to reward feedback interacts with phasic dopamine levels to determine the effectiveness of learning, particularly in dynamic feedback situations where frequency and magnitude of rewards need to be integrated over time to make optimal decisions. Using both simulations and behavioral data in humans, we evaluated how reduced sensitivity to losses, relative to gains, leads to suboptimal learning in the Iowa Gambling Task (IGT), a complex value-learning task. In the behavioral data, we tested individuals with a variant of the human dopamine receptor D2 (DRD2; -141C Ins/Del and Del/Del) gene that associates with lower levels of D2 receptor expression (N=119) and compared their performance to non-carrier controls (N=319). The magnitude of the reward response was measured by looking at ventral striatal (VS) reactivity to rewards in the Cards task using fMRI. DRD2 variant carriers had generally lower performance in the IGT than non-carriers, consistent with reduced sensitivity to losses. There was also a positive association between VS reactivity and performance in the IGT, however, we found no statistically significant difference in this effect between DRD2 carriers and non-carriers. Thus, while reduced D2 receptor expression was associated with less efficient learning in the IGT, we did not find evidence for the moderation of this effect by the magnitude of the reward response.

## Introduction

Consider the problem of choosing where to get your lunch: do you go with the food truck that always serves consistent mediocre food or the truck that sometimes serves amazing food, but at other times is simply unpalatable? Formally this represents a reinforcement learning problem (Sutton & Barto, 1998) with dynamic, or non-stationary, feedback schedules (Daw et al., 2006), which requires updating the estimated value of each action based on the gains (e.g., deliciousness) or losses (e.g., unpalatable) experienced in the past. From an algorithmic perspective, learning from these gains and losses happens in the form of temporal-difference (TD) learning (Sutton & Barto, 1998) that updates the expected value of any given action for any given state of the world. Over time this TD learning can lead to the optimal solution for determining action value, known as the Bellman solution (Bellman, 1956).

In the brain, TD learning is implemented, by phasic dopamine (DA) signals in cortico-basal ganglia-thalamic (CBGT) pathways (Fig. 1a). The CBGT pathways are organized as a set of computational loops, where each loop can be conceptually thought of as an independent decision (or action) channel (Mink, 1996; Bogacz & Gurney, 2007; Bogacz, 2007; A. Klaus et al., 2017). The goal of the CBGT loops is to integrate information from competing cortical sources to bias downstream selection systems towards one decision or another and then use feedback signals to promote learning that modifies this bias for future decisions (Mink, 1996). The canonical model of CBGT pathways relies on three dissociable control pathways: the direct (facilitation), indirect (suppression), and hyperdirect (braking) pathways. At any given moment, the instantaneous competition between the direct and indirect pathways reflects the strength of bias for a given decision (Dunovan & Verstynen, 2016; Bariselli et al., 2019; Mikhael & Bogacz, 2016). Reward feedback influences this biasing signal via phasic DA signaling (Schultz, 1998), which modifies the sensitivity of cortical signals on striatal spiny projection neurons. Specifically, the phasic DA signal is thought to reflect something akin to a reward prediction error (RPE; Schultz et al., 1992; Schultz et al., 1997). Positive RPEs, i.e., greater than expected gains, sensitize the D1-expressing cells of the direct pathway and depress the D2-expressing cells of the indirect pathway. Negative RPEs do the opposite, enhancing the sensitivity of the indirect pathway while depressing those of the direct pathway (Gurney et al., 2015). This opposing plasticity between the two pathways means that gains reinforce the appropriately selected action over less rewarding alternatives, while losses reduce the saliency of the selected action and allow for more competition between the action channels, pushing the network into a more exploratory state (Cools et al., 2009; Stauffer et al., 2014; Collins & Frank, 2015; Dunovan et al., 2019; Vich et al., 2020).

**Figure 1.**
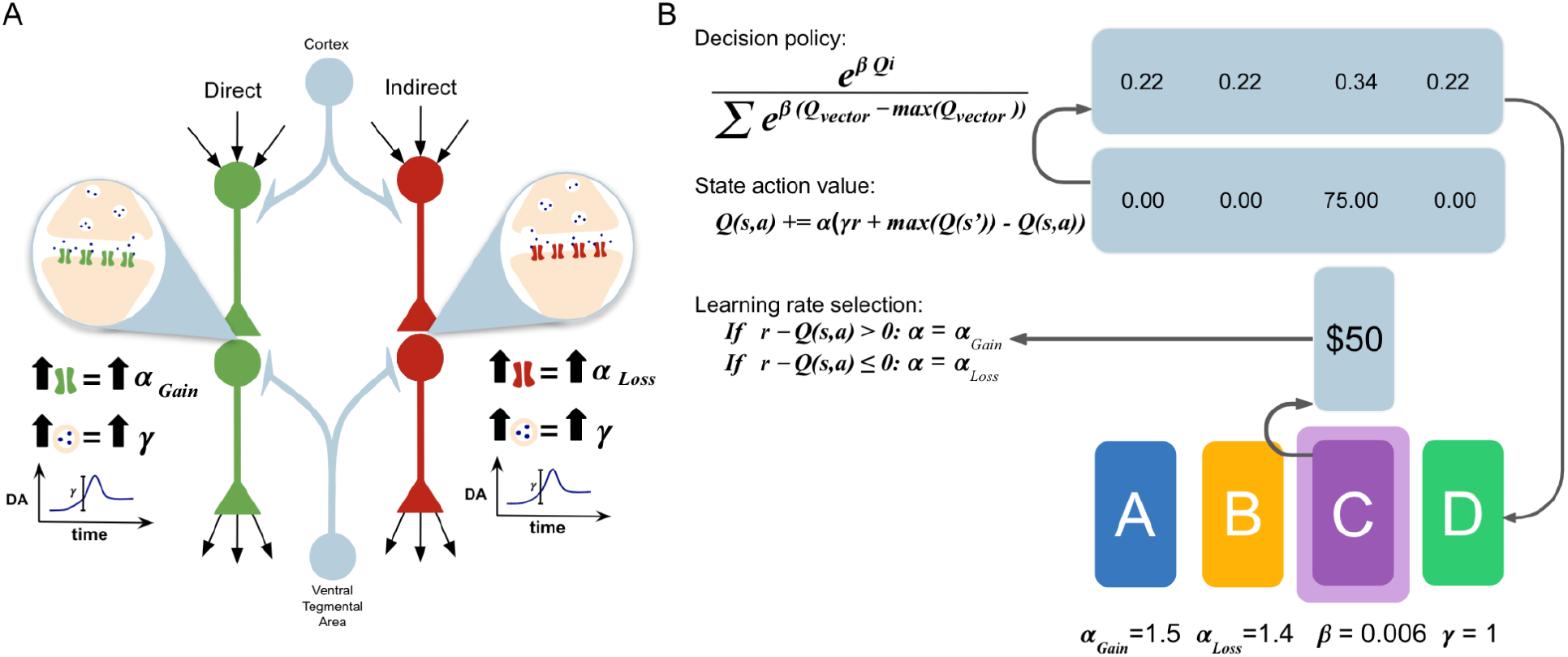
(**A**) Diagram of the two main pathways in the basal ganglia: direct facilitator pathway with D1 receptor neurons, and the indirect suppression pathway with D2 receptor neurons. (**B**) The Q-Learning Agent uses an error-driven learning algorithm coupled with the softmax exploration strategy to complete the Iowa Gambling Task. The learning paraments of the agent are abbreviated as: r = reward, RPE = r - Q(a), α_*Gain*_ = Learning Rate when RPE > 0, α_*Loss*_ = Learning Rate when RPE ≤ 0, β= Inverse Temperature (degree of randomness), γ = Reward strength (representative of dopamine reactivity).

There is another factor that influences how gains and losses impact learning: initial sensitivity to DA in the first place. Particular DRD2 gene polymorphism variants have been found to associate with functional modulation of dopamine receptor expression throughout the brain, including the striatum (Zhang et al., 2007). These variants have been shown to have a detectable influence on behavior, particularly learning. For example, individuals with these variants show blunted probabilistic learning in simple bandit-like tasks with strict probabilistic feedback (Frank et al., 2007; Frank & Hutchison, 2009; Gorwood et al., 2012; Jocham et al., 2009; Klein et al., 2007; K. Klaus et al., 2019; Foll et al., 2009).

Here we investigate how asymmetries in feedback sensitivity, driven by inherited differences in D2 receptor expression, might interact with phasic dopamine signals when learning to make value-based decisions in an environment where reward feedback is dynamic and, in some cases, deceptive. We hypothesize that the presence of the DRD2 polymorphism variant that associates with lower striatal D2 receptor density interacts with the magnitude of the evoked reward response measured using fMRI, to impact sensitivity to losses during learning in the Iowa Gambling Task.

## Materials & Methods

### Participants

We used an already collected sample of neurologically healthy adults from southwestern Pennsylvania taken from the University of Pittsburgh’s Adult Health And Behavior project, Phase II (AHAB-II). The sample consisted of 438 participants (228 females, 210 males, 81.7% White, non-Hispanic) between the ages of 30 and 54 years (M=42.67, SD=7.36). Every participant had their blood drawn and genotyped for the presence of DRD2 −141C Ins/Ins, Ins/Del, or Del/Del variants (see Lerman et al., 2005 for a detailed description of this method). Carriers were defined as having at least one deletion allele (i.e. DRD2 −141C Ins/Del, or Del/Del variants). The sample consisted of 119 carriers (55 male, 97 Ins/Del, 22 Del/Del), and 319 non-carriers (155 male). These participants were part of a larger project that included the completion of many other tasks, some of which were completed within the magnetic resonance imaging (MRI) scanner. This research project was approved by the institutional review boards at both the University of Pittsburgh and Carnegie Mellon University.

### Ventral Striatal Reactivity Task

The Cards task was used to assess the reactivity of the ventral striatum (VS) to negative and positive feedback cues associated with monetary gain (Hariri et al., 2006; Gianaros et al., 2011). The task consisted of 45 trials, divided into 9 separate blocks with 5 trials each. Within each trial, the participant was shown a “?” in the center of the card for 3 s, which indicated that the participant needed to now guess whether the following card would be less than or greater than 5. Their choice was indicated by a button press. An index finger press signaled less than 5, and a middle finger press signaled greater than 5. After the guess was made, the participant was presented with a number the computer selected for 500 ms, and then feedback for their response for 500 ms. The feedback was either a green up arrow for positive feedback for a correct response, or a red down arrow for negative feedback for an incorrect response. The end of the trial was then signaled with a cross-hair presented for 1.5 s. The total length of a trial was 5.5 s.

Each of the blocks was one of 3 different conditions: win, loss, or control. In the win condition, there was an 80% positive feedback rate (4 out of 5 correct responses) and a 20% negative feedback rate (1 out of 5 incorrect responses). The opposite was true for the loss condition. In the control condition, instead of receiving feedback or being asked to guess, they were presented with an “x” for 3 s and then instructed to press with either their index or middle finger in response. After pressing, they were then presented with an “*” for 500 ms and then a yellow circle for 500 ms. The block type varied by presenting “Guess Number” for 3 s at the start of each block for the win and loss conditions or “Press Button” for the control condition. The length of the task in total was 350 s.

Participants were scanned on a 3 T Trio TIM whole-body scanner (Siemens, Erlangen, Germany) using a 12-channel phased-array head coil (FOV) = 200 × 200 mm, matrix = 64 × 64, repetition time (TR) = 2000 ms, echo time (TE) = 29 ms and flip angle (FA) = 90◦, for more information see (Verstynen et al., 2020). While in the MRI scanner, participants completed a computerized reward task paradigm (for preprocessing information see (Verstynen et al., 2020). After preprocessing, linear contrast images, reflecting relative BOLD signal changes (i.e. win blocks versus loss blocks), were estimated for each participant using general linear model (GLM) estimation. The mean BOLD contrast parameter estimates were extracted from a predefined VS ROI (Verstynen et al., 2020; Gianaros et al., 2011). For more information on the estimation process and creating the a priori ROI mask, see (Verstynen et al., 2020), and Gianaros et al., 2011, respectively.

### Iowa Gambling Task

To measure decision-making in a dynamic and deceptive feedback environment, participants completed a computerized version of the Iowa Gambling Task (IGT). The IGT is a common task for assessing executive function in healthy and clinical populations (Buelow & Suhr, 2009). In the IGT, participants are asked to select a card from any of the four decks presented with a varying amount of reward or punishment (Bechara et al., 1994). The participants specifically select one card at a time from any of the 4 decks for a total of 100 card selections. The exact value and order of each of the cards within the 4 decks has been predetermined by the experimenters without the participant’s knowledge. The participants receive a loan of $2000 and are instructed that the goal of the task is to maximize profits. They are allowed to switch between any of the decks at any time and as often as they wished. The participants are not aware of any of the deck specifications and are only informed that each deck was different. With each selection from Decks A or B (the “disadvantageous decks”), participants have a net loss of money. With each selection from Decks C or D (the “advantageous decks”), participants have a net gain of money or a net of zero, respectively. The amount of reward or punishment varies between decks and the position within a deck. Deck A and Deck B both have the same amount of overall net loss. However, in Deck A the reward is less frequent and higher in magnitude, while in Deck B the reward is more frequent and higher in magnitude. Similarly, Deck C and Deck D have the same overall net gain. In Deck C the reward is less frequent and lower in magnitude, while in Deck D the reward is more frequent and higher in magnitude. From the selections made by the participants, their overall Payoff score (i.e. *payoff* = (*C* + *D*) − (*A* + *B*)) was calculated.

### Statistical Analysis

Group differences in VS reactivity and Payoff score were first evaluated using t-tests. Follow-up regression models, measured how VS reactivity, carrier status, and their interaction, associated with the Payoff score. Of particular interest are any potential race effects on gene-behavior associations. However, the non-white portion of the sample in the dataset was small (18.26%), yet made up a significant portion of the carriers (43.70%), making any independent racial group analyses severely underpowered in this data set. Nonetheless, we used model comparison procedures to determine whether age, self-reported gender, and racial identity needed to be included in the final regression model. The model with the control factors (AIC = 4191, BIC = 4220) was found to explain a negligible amount of additional information compared to the simpler model (AIC = 4211, BIC = 4228, Bayes factor = 18.370). Thus, for our final regression model, we opted to not include age, gender, and racial identity as control factors.

### Reinforcement Learning Agent

In order to simulate how different reward reactivities and learning rates impacted decision-making, we simulated IGT performance using a standard Q-learning agent with *a* softmax decision policy (Barto & Sutton, 1995). Q-learning is a specific form of TD learning where updates influence the subjective value of individual actions, as opposed to individual environmental states. The model equations are shown in Figure 1b. Briefly, on any given trial, the model selects one of four decks using a softmax decision policy (*P*). The inverse temperature parameter (β), which determines the greediness or randomness of the decision policy, was set to 0.006 for the final model, producing a moderately exploratory agent. After selection, a reward (*r*) is generated according to the feedback schedule of the IGT. Reactivity to reward is approximated by the scaling term γ, which is directly applied to *r*. The difference between the experienced reward, γ*r*, and the expected reward, *Q*(*a*), produces the reward prediction error (RPE) that is used to update *Q*(*a*) on the next trial. The learning rate (α), determines how much the RPE influences the update of *Q*(*a*). To approximate asymmetries in learning, we put a contingency on α. If the RPE is less than 0, then α = α_*Gain*_. Otherwise, for positive RPEs, α = α_*Loss*_.

For each simulation run, an agent was generated with a specific set of values for β, γ, α_*Gain*_, and α_*Loss*_. The agent completed 100 trials of the IGT with a predetermined deck, where the optimal deck is C. The trial-wise selection of decks across trials was used to calculate Payoff scores in the same way as estimated for human participants.

## Results

### Model simulations

In order to understand how asymmetries in learning from gains versus losses can impact the efficiency of decision-making in the IGT, we used a standard Q-learning agent (see Materials & Methods), where we varied the parameters specified in the decision and learning processes. The parameters of this model approximate the differences in decision-making between individuals in the IGT, specifically more exploratory or exploitative decision policies that impact long-term payoffs or losses. These parameters include sensitivity to positive RPEs (α_*Gain*_), sensitivity to negative RPE (α_*Loss*_), “greediness” of the decision policy (β), and overall sensitivity to reward (γ; see Materials & Methods and Figure 1B). The difference between the learning on gains versus losses can be reflected as asymmetrical such as when learning is stronger for cases where the reward value is greater than the expected value (RPE > 0; gains) and weaker when for when the reward value is less than the expected value (RPE ≤ 0; loss).

Figure 2 shows an example run of one of the agents. We see that over time the deck selections for this agent become more strategic, with a preference for decks C and D, with optimal deck C chosen the majority of the time (Figure 2A). This preference is reflected as an increase in state-action value (Q) for deck C in later trials (Figure 2B), which increases the probability that this deck will be selected over the others (Figure 2C). As a result, the optimal choices made by the agent increase over time as it effectively uses and learns from feedback, as seen by the percentage of choosing optimal deck C being above chance consistently after about 40 trials (Figure 2D).

**Figure 2.**
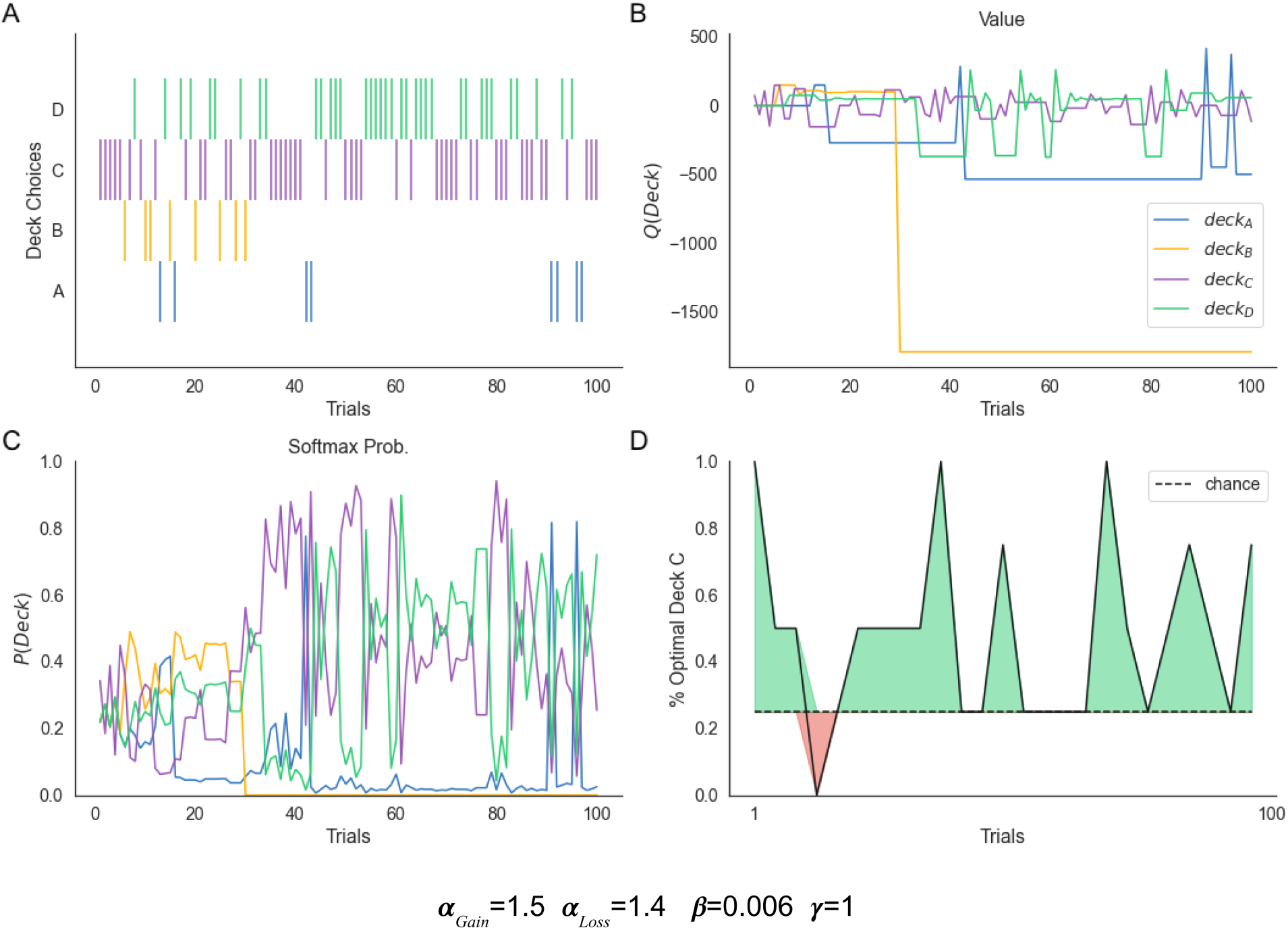
The parameters the agent was given are listed at the bottom of the figure. The agent completed 100 trials or card selections over decks A-D in which the deck choices were tracked in a raster plot (**A**), the predicted deck values were tracked in a line plot (**B**), the probability of selecting each deck was tracked in a line plot (**C**), and the percent optimal deck C is chosen is shown in the area plot (**D**).

In order to illustrate how the relative ratio of α_*Gain*_ and α_*Loss*_ impacts decision effectiveness (i.e., Payoff score), we ran a series of agents with different learning rate asymmetries and sensitivities to reward (Figure 3A-C). As expected, the heat maps in Figure 3A-C show that below a relatively low ratio of α_*Loss*_ to α_*Gain*_, the average Payoff score is negative. Payoff scores were greatest when α_*Loss*_ reached a level that allowed for the agents to learn from their mistakes, at around 0.2 to 0.3, depending on the overall sensitivity to rewards. This relationship between the ratio of α_*Loss*_ to α_*Gain*_ and performance in the task was amplified with an increase in sensitivity to rewards, γ (Figure 3D-F).

**Figure 3.**
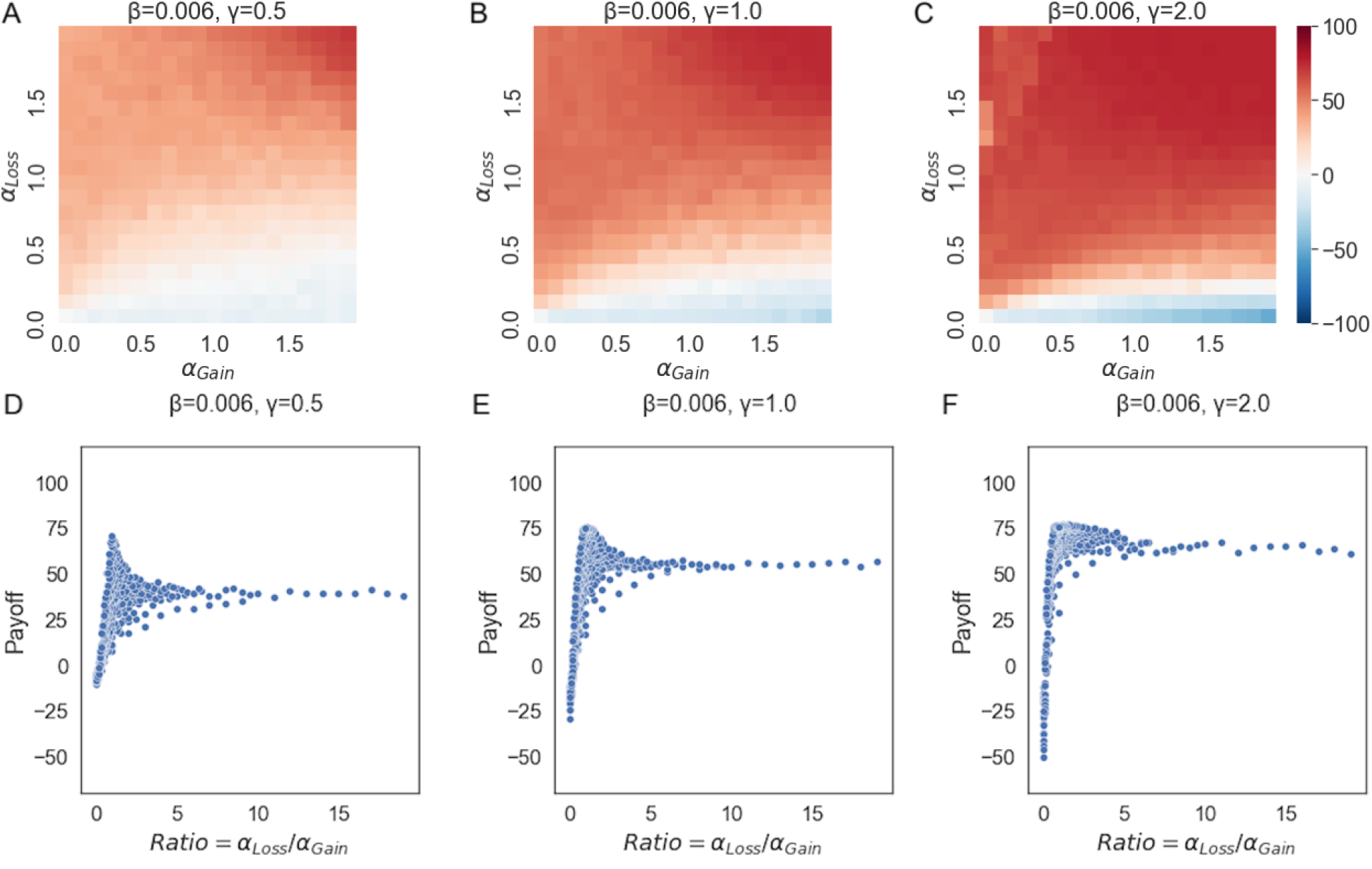
(**A-C**) Payoff scores were plotted in the heatmap for agents with alpha gain and alpha loss from 0-2, β of 0.006, and gamma of 0.5, 1, and 2. (**D-F**) Payoff scores were plotted against alpha ratio (alpha loss/alpha gain) for β of 0.006, and γ of 0.5, 1, and 2.

These simulations predict that in individuals with reduced learning from losses (i.e., negative RPEs) the overall Payoff scores in the IGT should be lower. Thus, we expect a main effect of the group on Payoff scores. If the groups also vary in sensitivity to rewards, this should also result in an interaction between the group and independent measures of reward sensitivity.

### Empirical Data

Our model simulations show that reduced sensitivity to negative feedback signals should reduce Payoff scores, and this effect should be scaled by how sensitive, or reactive, an individual is to rewards overall. To empirically test this we first looked at overall group differences in both Payoff scores and VS reactivity, in our sample of human participants. Consistent with our model, Payoff, the measure of effective use of feedback to make decisions, was significantly lower in the DRD2 carrier group than in non-carrier controls (Figure 4A; t[436] = −3.230, p = 0.001).

**Figure 4.**
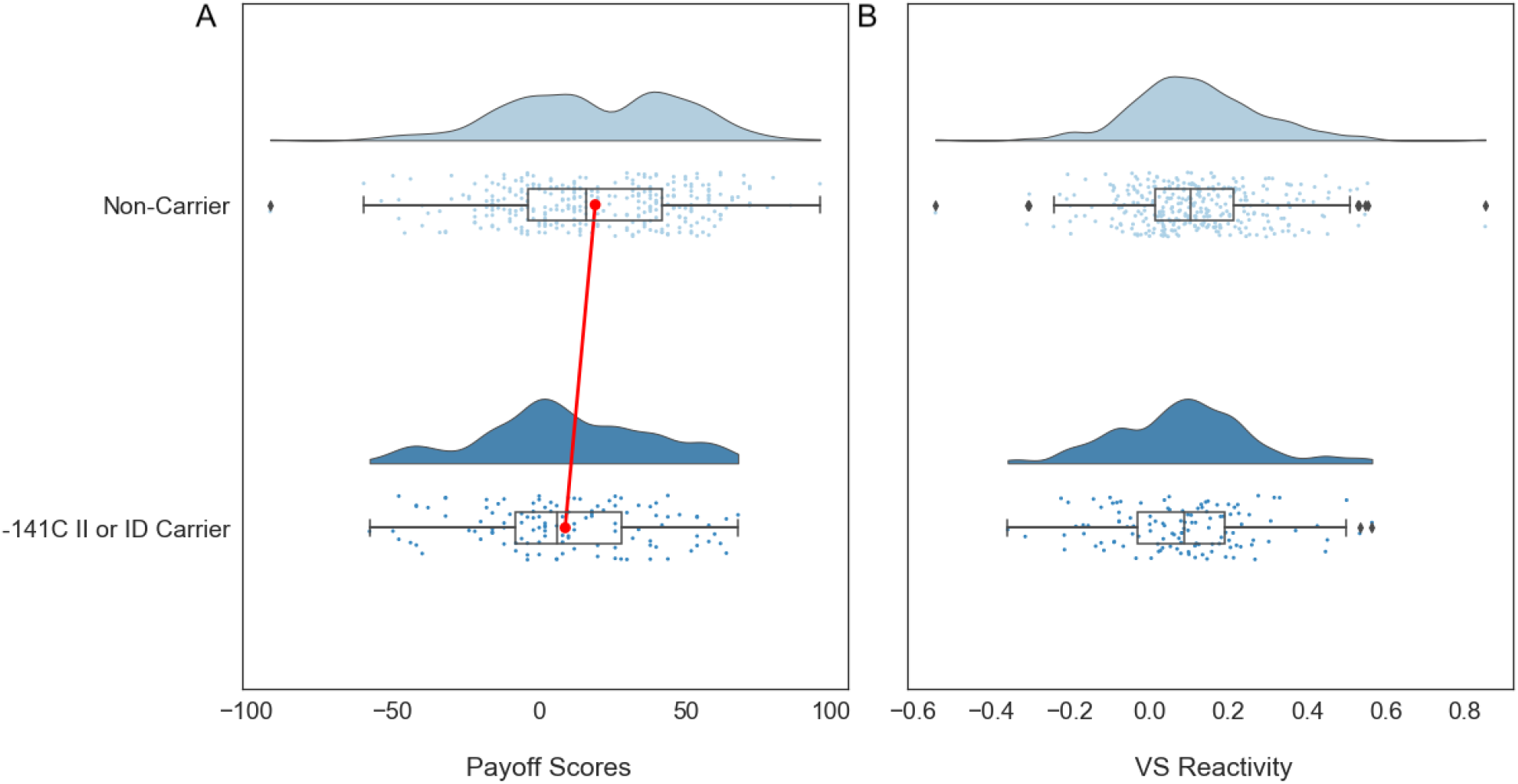
The distributions of measures for (**A**) IGT Payoff scores and (**B**) VS reactivity for each DRD2 group for non-carriers and carriers were plotted for comparison. The red line between mean Payoff scores between non-carriers and carriers demonstrates a significant difference in means.

Overall the non-carrier group had a mean Payoff score of 19.16, with a slightly bimodal distribution, whereas the carrier group had a more unimodal distribution, with a mean of 8.89. In contrast, VS reactivity was statistically equivalent between both the non-carrier and carrier groups, centered just above 0.0 for both (Figure 4A; t[436] = −1.771, p = 0.077). Thus, individuals with expected lower D2 receptor density perform worse in the IGT than controls but don’t show reliable differences in VS responses to rewards.

The model predicted that the impact of carrier status (approximating α_*Loss*_) on Payoff scores should be stronger in individuals with stronger VS reactivity (approximating γ). Given the lack of group differences in VS reactivity, we did not expect a reliable group-by-VS reactivity interaction on Payoff scores, even though one could still be possible. Figure 5A shows that both carriers and non-carriers have positive-trending slopes for the association between VS reactivity and Payoff score, consistent with our model (see above), with carriers having a slightly shallower trendline compared to the non-carriers. Indeed, this lack of an interaction effect is born out in a linear regression analysis shown in Table 1. We see that carrier status has a statistically significant, negative effect on Payoff score, replicating the t-test results. In contrast, an individual’s VS reactivity score correlated with an overall higher average Payoff score. However, the interaction term between the group and VS reactivity was not statistically significant. Thus, we do not see that higher VS reactivity amplifies the effect of group status on Payoff. Though this may not be surprising given the lack of overall group differences in VS reactivity.

**Table 1.**
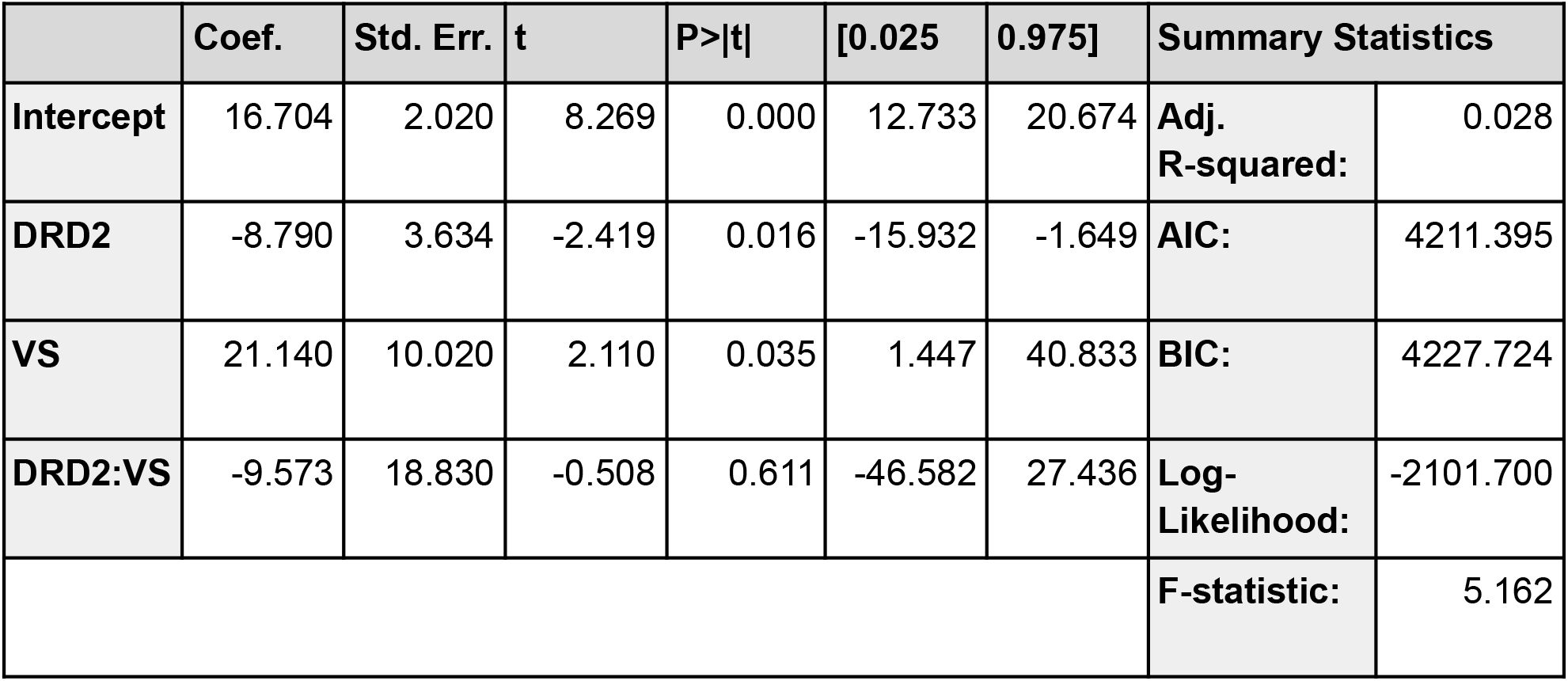
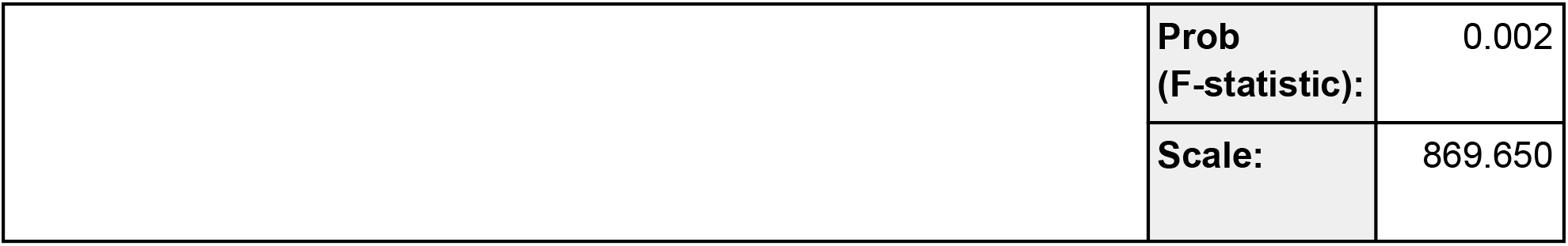
The linear regression model for Payoff was composed of only the main effect variables, the DRD2 group assignment, VS Reactivity, and their interaction. The summary results for the linear regression model were also included such as the AIC and BIC scores.

|  | Coef. | Std. Err. | t | P>\|t\| | [0.025 | 0.975] | Summary Statistics |  |
| --- | --- | --- | --- | --- | --- | --- | --- | --- |
| **Intercept** | 16.704 | 2.020 | 8.269 | 0.000 | 12.733 | 20.674 | **Adj. R-squared:** | 0.028 |
| **DRD2** | -8.790 | 3.634 | -2.419 | 0.016 | -15.932 | -1.649 | **AIC:** | 4211.395 |
| **VS** | 21.140 | 10.020 | 2.110 | 0.035 | 1.447 | 40.833 | **BIC:** | 4227.724 |
| **DRD2:VS** | -9.573 | 18.830 | -0.508 | 0.611 | -46.582 | 27.436 | **Log-Likelihood:** | -2101.700 |
|  |  |  |  |  |  |  | **F-statistic:** | 5.162 |

|  | Coef. | Std. Err. | t | P>\|t\| | [0.025 | 0.975] | Summary Statistics |  |
| --- | --- | --- | --- | --- | --- | --- | --- | --- |
| Intercept | 16.704 | 2.020 | 8.269 | 0.000 | 12.733 | 20.674 | Adj.<br>R-squared: | 0.028 |
| DRD2 | -8.790 | 3.634 | -2.419 | 0.016 | -15.932 | -1.649 | AIC: | 4211.395 |
| VS | 21.140 | 10.020 | 2.110 | 0.035 | 1.447 | 40.833 | BIC: | 4227.724 |
| DRD2:VS | -9.573 | 18.830 | -0.508 | 0.611 | -46.582 | 27.436 | Log-Likelihood: | -2101.700 |
|  |  |  |  |  |  |  | F-statistic: | 5.162 |
|  |  |  |  |  |  |  | Prob<br>(F-statistic): | 0.002 |
|  |  |  |  |  |  |  | Scale: | 869.650 |

**Figure 5.**
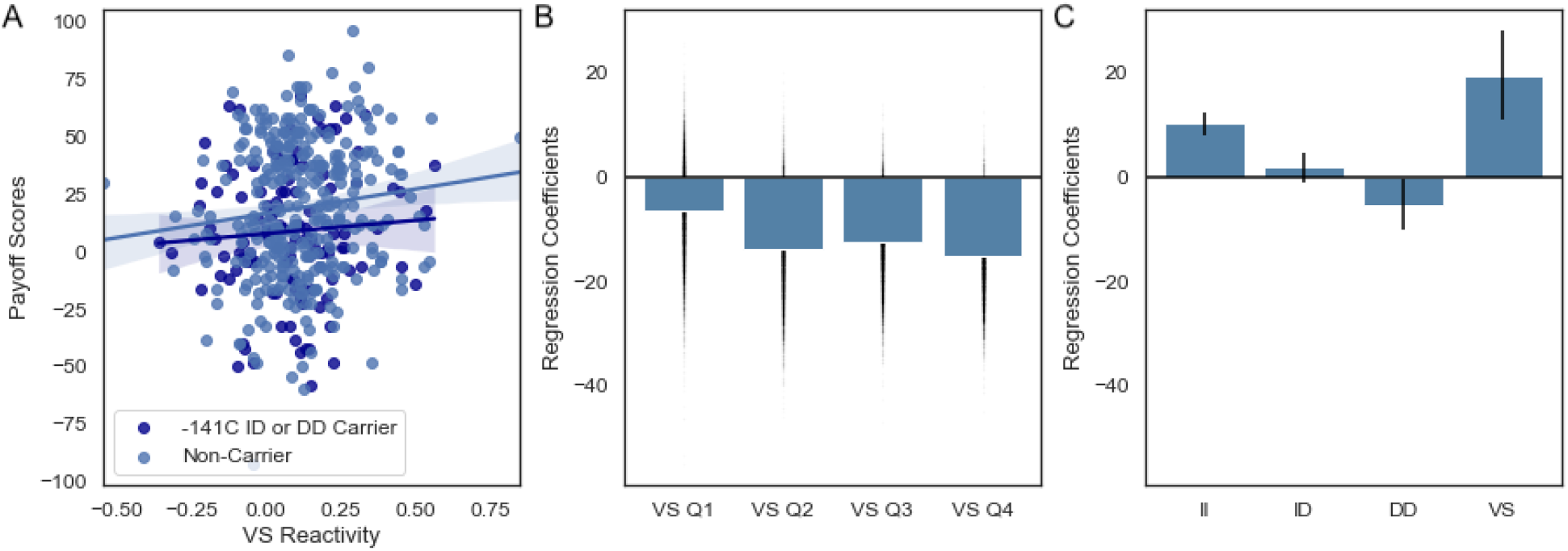
(**A**) Linear regression model was used for the visualization of the relationships between Payoff (P) with VS reactivity for both groups. (**B**) Bootstrapped regression model coefficients for carrier status on Payoff score, binned by VS reactivity quartiles. (**C**) Regression model coefficients for Payoff score versus DRD2 carrier status −141C Ins/Ins (II), Ins/Del (ID), Del/Del (DD), and VS reactivity (VS). Error bars reflect the standard error of the mean.

One assumption in our model simulations is that the effect of sensitivity to rewards on the expression of asymmetric learning is monotonic and linear. It is possible that this is not a valid assumption. Therefore, we took a closer look at the interaction between VS reactivity and DRD2 carrier status by binning participants according to VS reactivity quartiles, measuring the main effect of the group in each quartile separately. Three out of the four binned groups showed a negative effect of group status on Payoff scores, consistent with our overall effects. However, this did not increase as VS reactivity increased (Figure 5B). This enticing finding suggests that how sensitivity to reward interacts with asymmetries in learning may be more complicated than assumed in our simple reinforcement learning model. Though we cannot make any strong conclusions on this relationship due to the high variability across quantiles.

In addition to the binary classification of carriers and non-carriers, we also tested the linear regression model that included all 3 different DRD2 polymorphism variants, −141C Ins/Ins, Ins/Del, and Del/Del. Breaking down the main effect model into these 3 subgroups we see a tiering effect where an increase in Del alleles is associated with a decrease in Payoff scores. The regression coefficients of this model for the different groups, as well as VS reactivity, are shown in Figure 5C. It is important to note that the sample sizes of each respective DRD2 polymorphism group (Ins/Del N=97, Del/Del N=22) leave us with low statistical power to look at more complex models. However, the main effects analysis clearly discerns a reliable group effect.

## Discussion

Here we investigated how differences in striatal D2 receptor expression could interact with sensitivity to rewards to impact feedback-based learning in situations where reward feedback is dynamic and deceptive. Using a simple reinforcement learning model, we first showed how asymmetries in learning on gains vs. losses should impact performance in the IGT, and how this can be moderated by reward reactivity. Our experimental data in human participants support part of our computationally driven hypothesis, in that individuals with a genetic predisposition for lower D2 receptor expression performed worse on the IGT than controls. However, this effect was not scaled by individual differences in reward reactivity as predicted by our model. Nonetheless, our results further bolster the observation from both the computational modeling and behavioral literature that sensitivities to losses are a critical component in effective long-term learning, particularly in dynamic and deceptive feedback environments.

These findings align with the prior literature demonstrating that genetic variants influencing D2 receptor expression also impact goal-directed behavior. For example, Zhang et al. 2007 looked at two SNPs that regulate the D2 receptor and found that carriers of the SNPs, which downregulate D2 receptor expression as in our study, showed altered striatal responses during an N-Back working memory task (Zhang et al., 2007). In contrast, we did not see that striatal responses in the functional imaging task differed between carrier groups. This may simply be due to the difference in the cognitive process being measured in the scanner (e.g., working memory vs. reward reactivity). Although the BOLD responses during the Cards task, used as a measure of VS reward reactivity, do not directly measure DA neurotransmission, the relative pattern has been found to be consistent with in vivo human striatal DA synthesis as measured through FDOPA in positron emission tomography (Siessmeier et al., 2006). Whether this evoked response represents a good proxy for the true phasic DA response requires further testing.

Along these same lines, Klein et al. showed that individuals with the TaqA1 polymorphism variant, which is also believed to reduce D2 receptor expression, showed reduced sensitivity to errors in a probabilistic avoidance task (Klein et al., 2007). This effect has been replicated multiple times (Frank et al., 2007; Frank & Hutchison, 2009). Our work extends this by showing how this reduced sensitivity to negative feedback, as a result of genetic predispositions for D2 receptor expression, impacts the efficiency of using feedback in more complex reinforcement scenarios where frequency and magnitude of gains or losses need to be integrated over time in order to make an optimal decision. Later, Jocham et al. (Jocham et al., 2009) showed that TaqA1 polymorphism variant carriers had deficits in reversal learning that consisted of a decreased ability to sustain the newly rewarded response after a reversal and a decreased tendency to stick to the rewarding response in general (Jocham et al., 2009). Taken together with our current work, these findings provide clear evidence that a reduction in D2 efficiency or expression can lead to deficiencies in the integration of feedback from positive and negative signals.

Understanding how our findings relate to the broader behavioral genetics literature on the influence of D2 pathways in cognition should be tempered by the heterogeneity of genetic polymorphisms on the underlying dopamine pathways. The TaqA1 (SNP ID: rs1800497), C957T (SNP ID: rs6277), and the −141C Ins/Del (SNP ID: rs1799732) polymorphism variants are some of the most well understood genetic factors impacting the D2 pathway, although there are many others being discovered and studied (Foll et al., 2009; Gorwood et al., 2012). Both the −141C Ins/Del and TaqA1 polymorphism variants are believed to primarily impact dopaminergic signaling by lowering the D2 receptor density in the striatum (Jönsson et al., 1999; Pohjalainen et al., 1998), whereas the C957T polymorphism is believed to be impacting dopamine D2 receptor availability by affecting the receptor affinity to dopamine (M. M. Hirvonen, Laakso, et al., 2009; M. M. Hirvonen, Lumme, et al., 2009; Smith et al., 2017). Furthermore, the −141C Ins/Del and C957T polymorphisms are believed to be directly on the DRD2 gene (Arinami et al., 1997; M. Hirvonen et al., 2004), whereas the TaqA1 polymorphism is located downstream of the DRD2 gene in the ankyrin repeat and kinase domain containing 1 (ANKK1) gene (Neville et al., 2004). Trying to integrate findings across these disparate genetic markers in order to come to a mechanistic understanding of how dopamine, particularly D2, pathways influence behavior requires a careful accounting of the different influences these mutations have on the underlying neural circuitry.

It is also worth noting that while our results and prior work suggest that mutations impacting D2 pathways impact high-level decisions, not all evidence points in this direction. A recent meta-analysis by Klaus et al. was not able to establish significant associations between the TaqA1 and C957T polymorphism variants and any of the executive function domains tested, which included a variety of batteries measuring working memory, response inhibition, and cognitive flexibility (K. Klaus et al., 2019). The results of this systematic review suggest that the presence of TaqA1 and C957T polymorphism variants and their impact in dopamine D2 receptors signaling may have a limited effect on high-level executive function. Although, it is also possible that the neuropsychological batteries covered in the review by Klaus and colleagues may not be sensitive to subtle variation in cognitive ability that may be driven by differences in DA receptor expression.

So where do the present results, and the broader literature, leave us in understanding the role of different DA systems during learning, particularly in contexts where feedback is dynamic and deceptive? Of primary importance for future work determining the mechanism by which sensitivity to feedback signals interacts might reward reactivity. Our reinforcement learning model, as well as general intuition, shows clearly that these two factors should interact, yet we failed to find this in our data (but see Verstynen et al., 2020). One possibility could be the need to find a better or more specific, marker of phasic dopamine responses. This would likely require moving to more invasive methods, likely in non-human model populations. Another possibility is that feedback signal sensitivity may have a non-linear interaction with VS reactivity, and a non-linear interaction model would need to be tested with a much larger sample size capable of discerning this interaction. Another open question is the role of D1 pathways in this learning process. Our model assumes that learning on gains and losses are equally important for effective long-term value learning, but our work, as well as prior work, only looks at the role of D2 pathways and learning from losses. Integrating findings across genetic markers for the different dopamine pathways would help to fully elucidate the nature of this process. Of course, these concerns can be tested both experimentally, and theoretically using biologically realistic models of corticostriatal plasticity during learning (e.g., Gurney et al., 2015; Vich et al., 2020). Working out these precise mechanisms of both learning on gains and losses as well as their possible interaction with reward reactivity is left to future work.

